# Proximity-based proteomics reveals the thylakoid lumen proteome in the cyanobacterium *Synechococcus* sp. PCC 7002

**DOI:** 10.1101/2020.04.07.030940

**Authors:** Kelsey K. Dahlgren, Colin Gates, Thomas Lee, Jeffrey C. Cameron

## Abstract

Cyanobacteria possess unique intracellular organization. Many proteomic studies have examined different features of cyanobacteria to learn about the structure-function relationships between the intracellular structures of cyanobacteria and their roles in cells. While these studies have made great progress in understanding cyanobacterial physiology, the previous fractionation methods used to purify cellular structures have limitations; specifically, certain regions of cells cannot be purified with existing fractionation methods. Proximity-based proteomics techniques were developed to overcome the limitations of biochemical fractionation for proteomics. Proximity-based proteomics relies on spatiotemporal protein labeling followed by mass spectrometry of the labeled proteins to determine the proteome of the region of interest. We have performed proximity-based proteomics in the cyanobacterium *Synechococcus sp.* PCC 7002 with the APEX2 enzyme, an engineered ascorbate peroxidase. We determined the proteome of the thylakoid lumen, a region of the cell that has remained challenging to study with existing methods, using a PsbU-APEX2 gene fusion. This study demonstrates the power of APEX2 as a tool to study the cell biology of intracellular features of cyanobacteria with enhanced spatiotemporal resolution.

## Introduction

The intracellular spatial organization of cyanobacteria is unique among prokaryotes. As Gramnegative bacteria, cyanobacteria possess the typical inner and outer membrane systems enclosing a cell wall comprised of peptidoglycan. However, most cyanobacterial species also possess thylakoid membranes, an extra set of intracellular membranes where photosynthesis occurs, as well as carboxysomes, proteinaceous organelles used for carbon fixation. The distinctive intracellular spatial organization and protein complexes found within cyanobacteria has drawn particular interest to the cell biology of these organisms. Furthermore, cyanobacteria can also be used as a model for plant chloroplasts, as they share features and have a common evolutionary ancestor. As a result, many proteomic studies of specific cyanobacterial structures, i.e. thylakoid membranes, have been performed (Agarwal et al., 2010; Fulda et al., 2000; Huang et al., 2002, 2004, 2006; Li et al., 2012; Liberton et al., 2016; Pisareva et al., 2007, 2011; Rajalahti et al., 2007; Srivastava et al., 2005; Wang et al., 2000; Zak et al., 2001; Zhang et al., 2009). These studies have made great progress towards understanding the physiology of cyanobacteria, but lack the spatial resolution necessary to resolve the composition of many intracellular compartments resistant to traditional biochemical fractionation and purification methodologies.

Previously, proteomic studies of cyanobacterial components were limited to fractionation and separation techniques which could introduce artifacts and result in ambiguous cellular localizations. Furthermore, existing techniques are impractical for non-membrane bound regions or the thylakoid lumen. However, a technique termed proximity-based proteomics was recently developed in mammalian cells to allow for proteomic analysis of cellular regions or protein interactomes that were unable to be purified using existing techniques (Kim and Roux, 2016). Proximity-based proteomics relies on targeting a specific enzyme to a region of interest as a protein fusion to a full-length protein or signal sequence. The enzyme then performs chemistry in live cells to label proteins within a small radius (10-20 nm) of itself (Rhee et al., 2013). After cell lysis, the labeled proteins can then be separated from unlabeled proteins and analyzed using mass spectrometry. Several proximity-based proteomics techniques exist, but the most common use enzymes that biotinylate proteins (Kim and Roux, 2016). We chose to use APEX2, an engineered ascorbate peroxidase that catalyzes a reaction between biotin-phenol (BP) and hydrogen peroxide (H_2_O_2_) to create a BP radical that covalently attaches to proteins (Hung et al., 2016; Lam et al., 2015) (Figure 1A). The reactivity and short half-life of biotin-phenol gives this technique a high spatial specificity. Furthermore, APEX2 has been shown to be catalytically active in multiple cellular compartments and exhibits a short (1 minute) labeling time, allowing for high temporal specificity (Hung et al., 2016; Lam et al., 2015).

**Figure 1.**
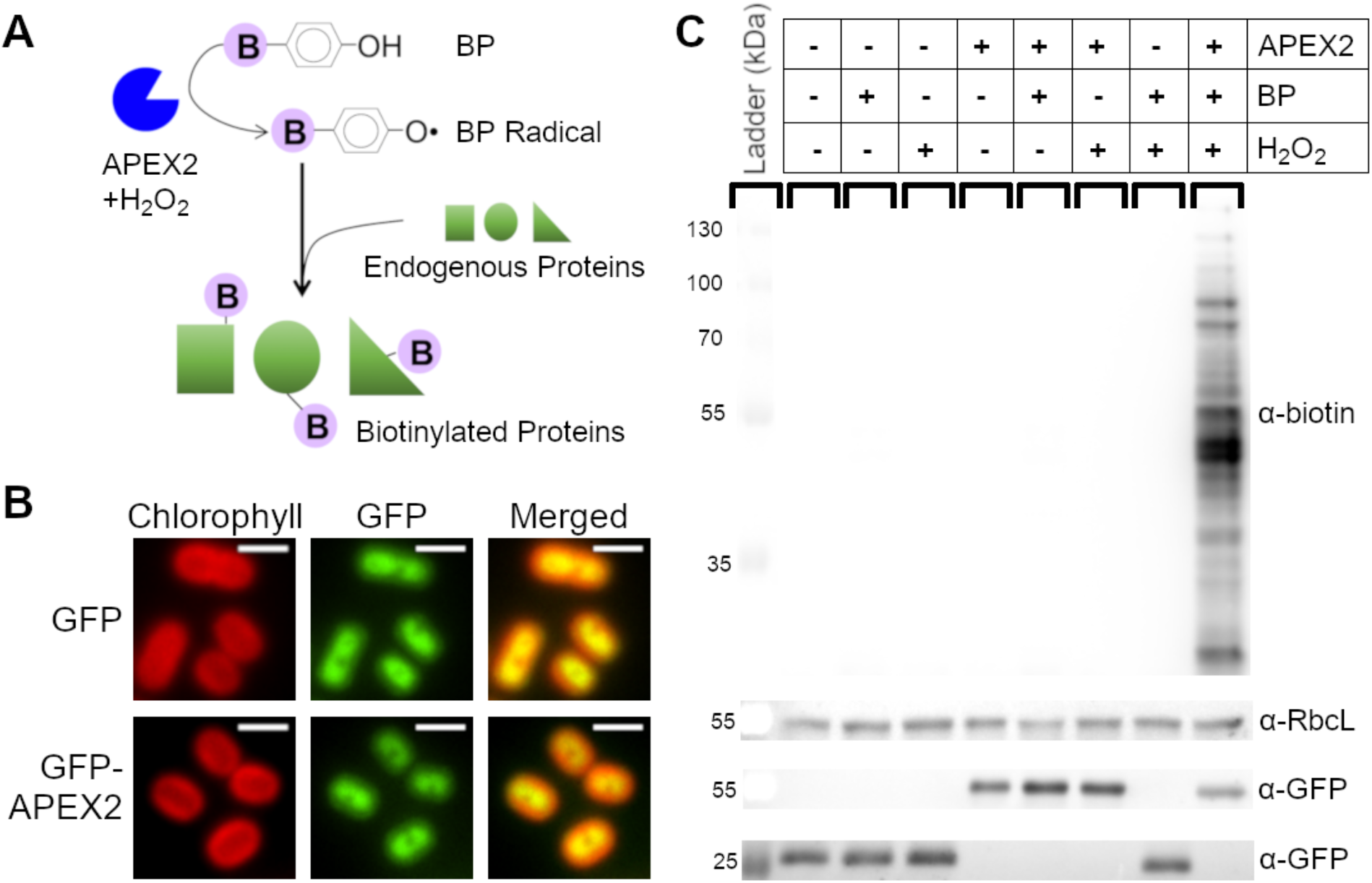
APEX2-dependent labeling specifically biotinylates proteins in PCC 7002. **(A)** APEX2 reacts with BP in the presence of H_2_O_2_ to produce a BP radical. Biotinylated proteins are generated when the BP radical reacts with peptides, forming a covalent bond. **(B)** Fluorescence microscopy imagines of cells expressing GFP and GFP-APEX2 (green). Scale bars are 2 μm. Chlorophyll channel (red) indicates thylakoid membrane. **(C)** 5 μg of protein from cells expressing either GFP or GFP-APEX2 was separated by SDS-PAGE and transferred to a membrane for immunoblot analysis using α-biotin to detect APEX2 activity. α-RbcL was used as a loading control and the same membrane was stripped and re-probed with α-GFP to check for expression of GFP (28 kDa) or GFP-APEX2 (54 kDa).

Here we demonstrate the feasibility and potential of proximity-based proteomics technique using APEX2 in *Synechococcus sp.* PCC 7002 (PCC 7002), a model cyanobacterium and promising chassis for biotechnological applications (Markley et al., 2015; Ruffing et al., 2016; Xu et al., 2011). To showcase the ability of APEX2 to interrogate regions of the cell where proteomics studies have not yet been possible due to limitations of existing biochemical methods, we targeted APEX2 to the thylakoid lumen by fusing it to PsbU, an extrinsic photosystem II (PSII) protein (Nishiyama et al., 1998), and identified the PsbU-associated proteome by mass spectrometry. Determining the thylakoid lumen proteome is vital for understanding the physiological roles of the thylakoid membrane system and the reactions of oxygenic photosynthesis.

## Results and Discussion

### Characterization of APEX2 labeling in PCC 7002

To determine if APEX2-dependent labeling of proteins was possible in cyanobacteria, GFP or GFP-APEX2 was incorporated into the genome of PCC 7002. Cytoplasmic localization of GFP and GFP-APEX2 was confirmed using fluorescence microscopy (Figure 1B). To perform APEX2-dependent biotinylation, cells were incubated with BP and then exposed to H_2_O_2_. After quenching the reaction, cells were lysed by bead beating and a streptavidin blot confirmed the ability of APEX2 to biotinylate proteins in PCC 7002 (Figure 1C). Biotin labeling only occurred in the presence of APEX2, BP, and H_2_O_2_ demonstrating reaction specificity in vivo. Furthermore, the rapid reaction enables precise temporal control of labeling.

### Purification of cytoplasmic APEX2-biotinylated proteins from PCC 7002

Proteins biotinylated in vivo were enriched for further analysis by affinity purification. APEX2-dependent biotinylation was performed in cells expressing GFP or GFP-APEX2 in the cytoplasm. Affinity purification of biotinylated proteins was performed by incubating cellular lysates with streptavidin-coated magnetic beads. The background level of biotinylation was very low as biotinylated protein was only detected in cells expressing GFP-APEX2, but not cells expressing GFP alone (Figure 2A-B). To confirm cytoplasmic APEX2 labels cytoplasmic proteins, immunoblots using antibodies against expected cytoplasmic proteins were performed (Figure 2C-D). Since the BP radical reacts with proteins within a 10-20 nm radius of its origin, APEX2 itself is expected to be biotinylated. Biotinylated GFP-APEX2 fusion protein was detected using a α-GFP antibody, confirming the expected self-reactivity (Figure 2C). Additionally, the large subunit of rubisco (ribulose-1,5-bisphosphate carboxylase/oxygenase), RbcL, an abundant cytoplasmic protein, was only enriched on beads cells incubated with cells expressing GFP-APEX2 as detected using a specific α-RbcL antibody (Figure 2D). The high molecular weight RbcL band in lysates is likely the result of higher-order complexes formed in vivo; RbcL is associated with large protein assemblies including the carboxysome in cyanobacteria (Cameron et al., 2013). Following the more stringent enrichment and elution process, these complexes have been disrupted and RbcL migrates as expected.

**Figure 2.**
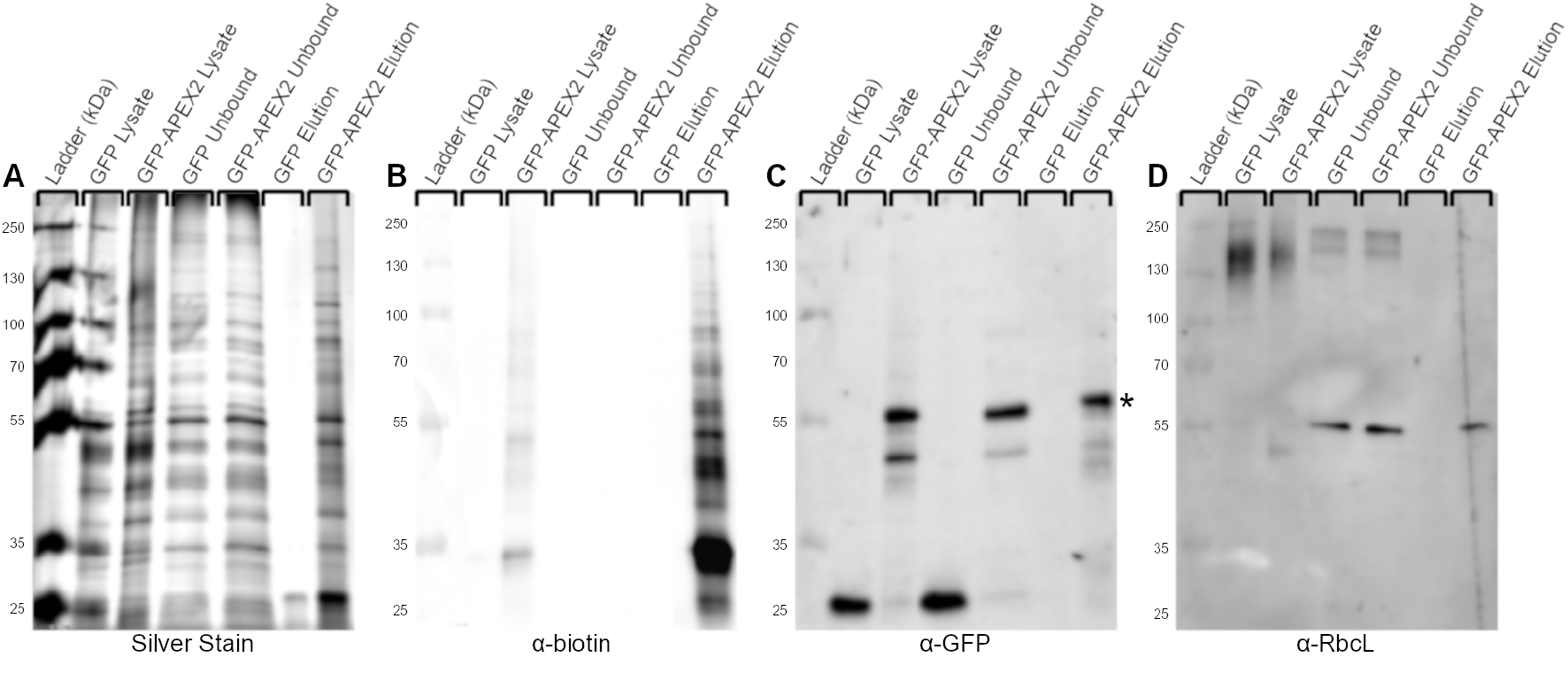
Enrichment of proteins biotinylated by cytoplasmic APEX2 in vivo. Cells expressing GFP or GFP-APEX2 were incubated with BP and exposed to H_2_O_2_. Biotinylated proteins were captured from cell lysates on streptavidin coated magnetic beads. Fractions from each enrichment step were separated by SDS-PAGE and then silver stained for contrast or transferred to a nitrocellulose membrane and probed with specific antibodies. **(A)** Silver stain of noted fractions from unlabeled (GFP) or labeled (GFP-APEX2) lysates. **(B)** Biotinylated proteins are only detected in fractions containing APEX2 and are enriched on streptavidin beads. **(C)** Expected self-labeling (biotinylation) of GFP-APEX2 (54 kDa, marked with *) is confirmed by immunoblotting against GFP. **(D)** RbcL (55 kDa), a cytoplasmic protein expected to be labeled by GFP-APEX2 was specifically captured on beads incubated with GFP-APEX2.

### PsbU-APEX2 and Cytoplasmic APEX2 label different sets of proteins

To demonstrate the ability of proximity-based proteomics to interrogate subcellular regions that have not been successfully purified using traditional purification methods, APEX2 was fused to an extrinsic subunit of photosystem II (PSII) in an effort to identify proteins in the thylakoid lumen. A PsbU-APEX2 gene fusion expressed from a neutral site in the chromosome under a constitutive promoter was used to localize APEX2 to the thylakoid lumen of PCC 7002. APEX2-dependent labeling and biotinylated protein purification was performed with PsbU-APEX2 and GFP-APEX2. A silver stain of purified biotinylated proteins from GFP-APEX2 and PsbU-APEX2 shows different banding patterns, suggesting that a different set of proteins is labeled by the different APEX2 fusions (Figure 3A). The thylakoid localization of PsbU-GFP was confirmed using fluorescence microscopy (Figure 3B). The localization of PsbU-GFP was used as a proxy for the localization of PsbU-APEX2, since GFP and APEX2 are both C-terminal tags of a similar size. To identify the proteins labeled by the different APEX2 fusion proteins, biotinylated proteins were purified from two independent samples of both PsbU-APEX2 labeled and GFP-APEX2 labeled cells, and the resulting peptides following tryptic digestion were separated and detected using LC-MS/MS. Label-free quantitative methods based on spectral counts were used to compare samples (Old et al., 2005). 99 proteins were identified exclusively in PsbU-APEX2 samples and 279 proteins identified exclusively in GFP-APEX2 samples. 437 proteins were identified in both PsbU-APEX2 and GFP-APEX2 samples (Figure 3C and Supplemental Table 1).

**Figure 3.**
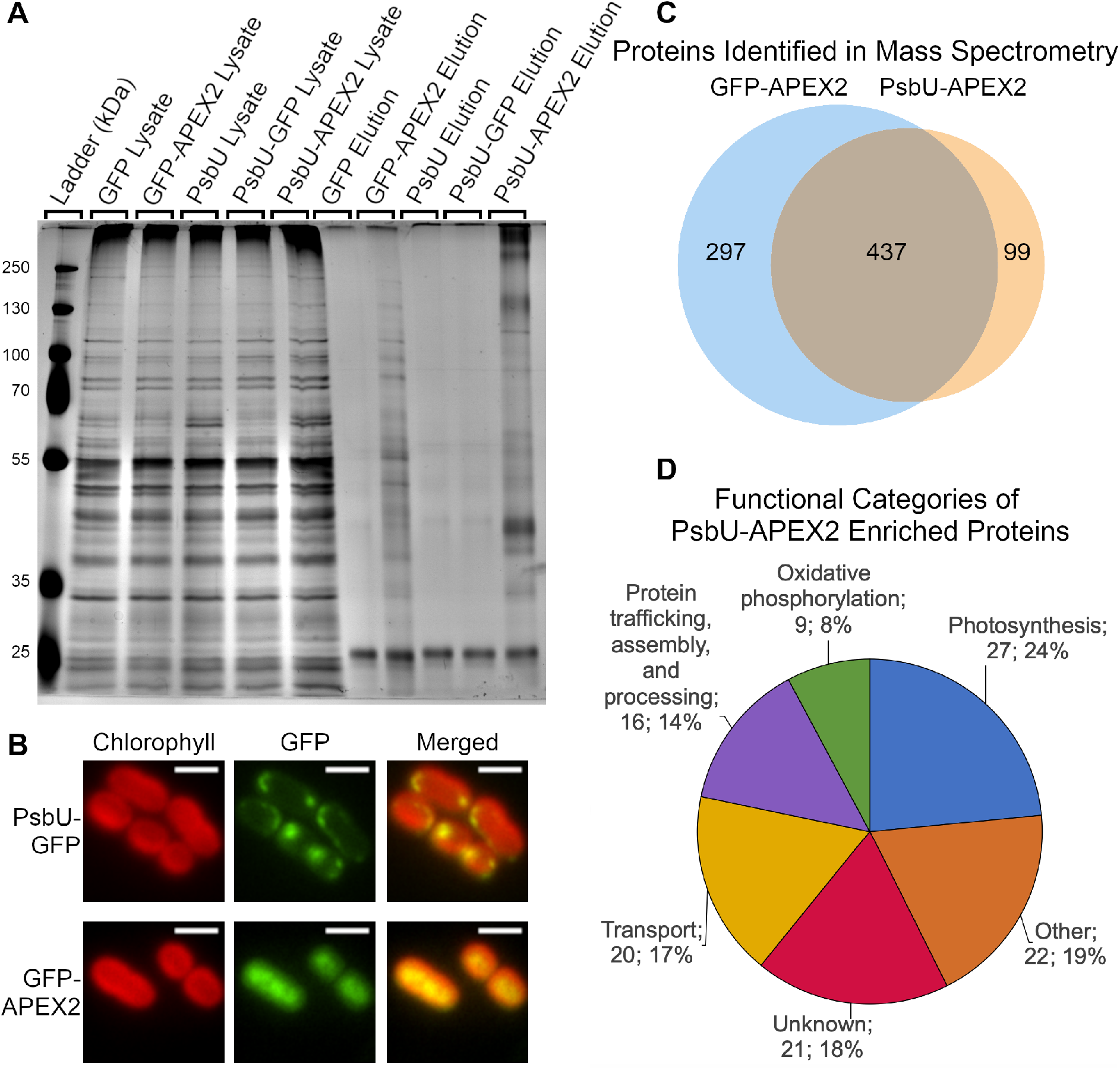
PsbU-APEX2 and Cytoplasmic APEX2 label different sets of proteins. **(A)** Silver stain of the biotinylated protein purification from PCC 7002 expressing GFP, GFP-APEX2, PsbU, PsbU-GFP, or PsbU-APEX2 cells exposed to BP and H_2_O_2_ **(B)** Localization of PsbU-GFP and GFP-APEX2 were visualized with fluorescence microscopy (Green). Chlorophyll channel indicates thylakoid membrane. Scale bars are 2 μm. **(C)** Biotinylated proteins from strains expressing GFP-APEX2 and PsbU-APEX2 identified by mass spectrometry (Also see Supplemental Table 1). **(D)** Functional categories of the proteins enriched in PsbU-APEX2 samples (number of proteins; percentage of 115 total proteins). The proteins used for this analysis are listed in Table 1. (Also see Supplemental Tables 1 and 2).

### Biotinylated proteins enriched in PsbU-APEX2 samples

Mass spectrometry data was further analyzed to determine what proteins were labeled by PsbU-APEX2. PsbU is a luminal extrinsic subunit of photosystem II and therefore the majority of proteins are expected to be localized to the thylakoid membrane or lumen. However, because PsbU-APEX2 is translated in the cytoplasm and then translocated to its final localization in the lumen, we also expected that a small population of PsbU-APEX2 could be present in the cytoplasm, resulting in labeling of cytoplasmic proteins. Therefore GFP-APEX2 was used as a control instead of a sample lacking APEX2/BP/H_2_O_2_, since it would control for the small cytoplasmic population of PsbU-APEX2 in addition to proteins nonspecifically bound to the streptavidin beads.

MaxQuant Label Free Quantitation (LFQ) intensities and normalized spectral counts were used to determine the identity of proteins specifically enriched with PsbU-APEX2 compared to the GFP-APEX2 control. Proteins were organized by descending enrichment (log_2_(PsbU-APEX2 LFQ intensity or spectral counts/GFP-APEX2 LFQ intensity or spectral counts). A true positive list, which includes proteins with evidence for thylakoid lumen or thylakoid membrane localization, was constructed from proteomic studies including localization data in *Synechocystis sp.* PCC 6803 (PCC 6803) (Agarwal et al., 2010; Fulda et al., 2000; Huang et al., 2002, 2004, 2006; Li et al., 2012; Liberton et al., 2016; Pisareva et al., 2011; Rajalahti et al., 2007, 2007; Srivastava et al., 2005; Wang et al., 2000; Zak et al., 2001; Zhang et al., 2009). A false positive list was constructed from proteins annotated as involved in DNA replication, transcription, or translation, because those proteins are expected to be localized to the cytoplasm (McClure et al., 2016). As expected, proteins from the true positive list have significantly higher enrichment values than proteins from the false positive list (Supplemental Figure 2). Using the true and false positive list, we identified a cutoff to discriminate between enriched proteins and those that bound to the beads non-specifically. This analysis was performed using enrichment values from both PsbU-APEX2 samples and was repeated using enrichment values calculated from normalized spectral counts and from LFQ intensities. The proteins above the cutoff in all four analyses are listed in Table 1. The major functions of enriched proteins are shown in Figure 3D.

**Table 1:**
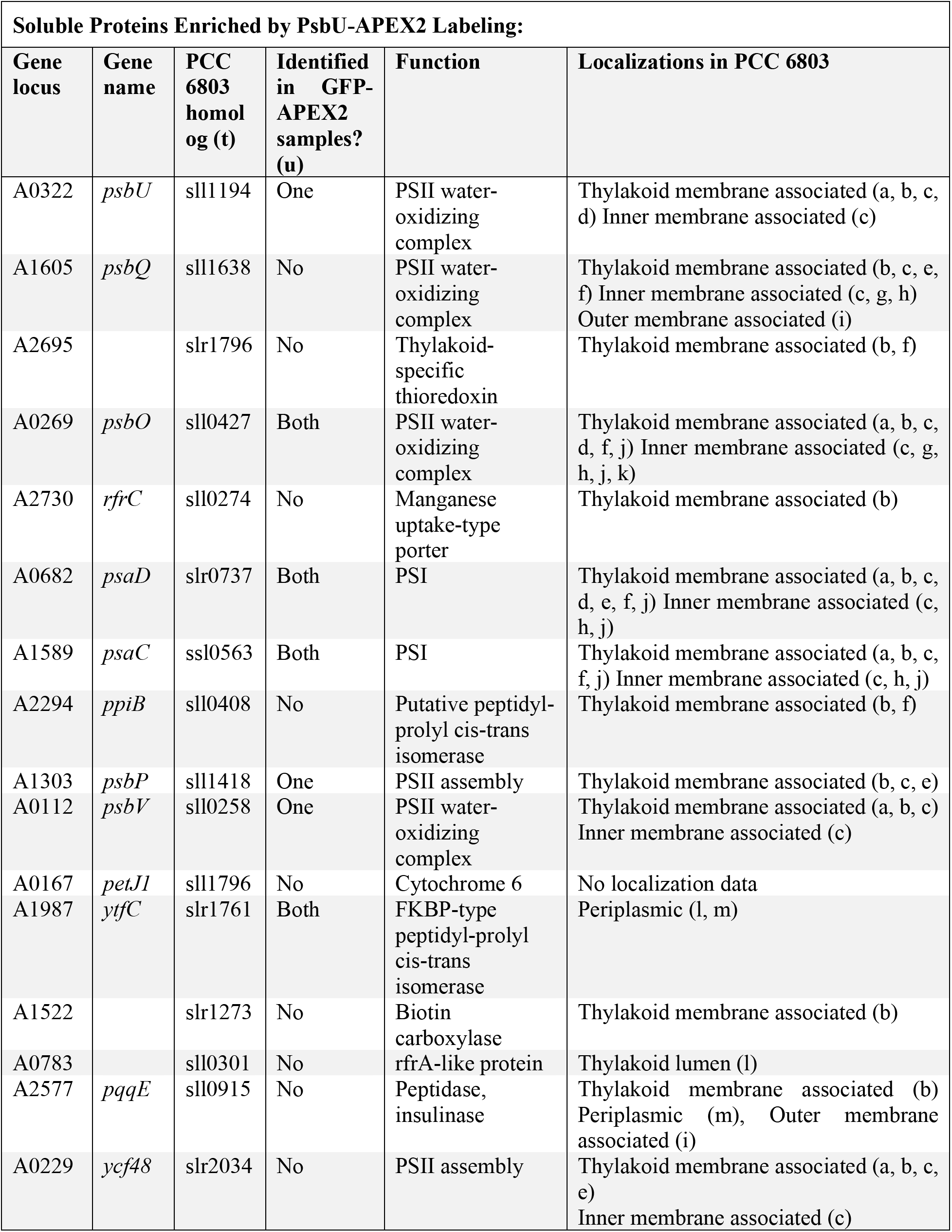
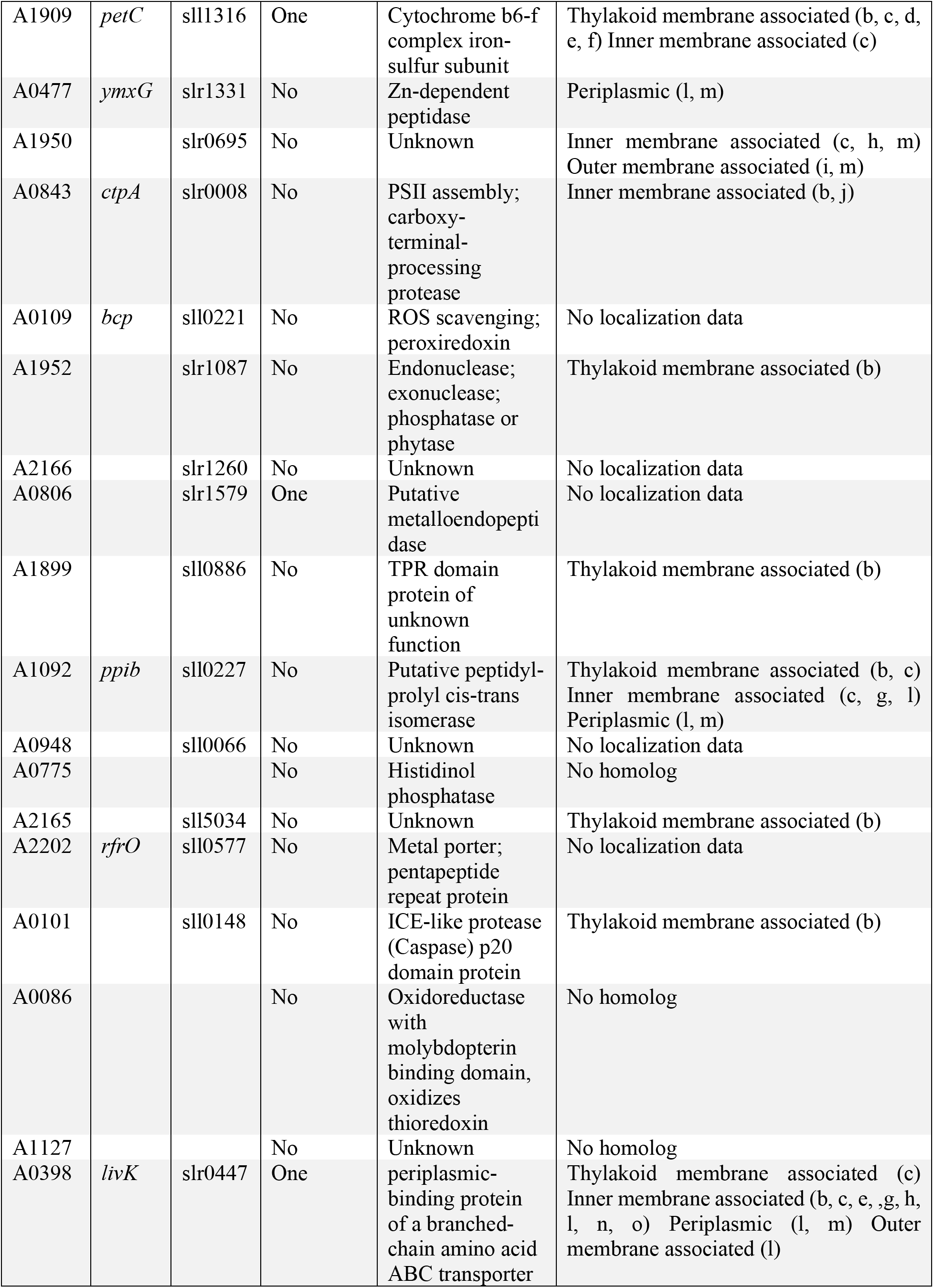
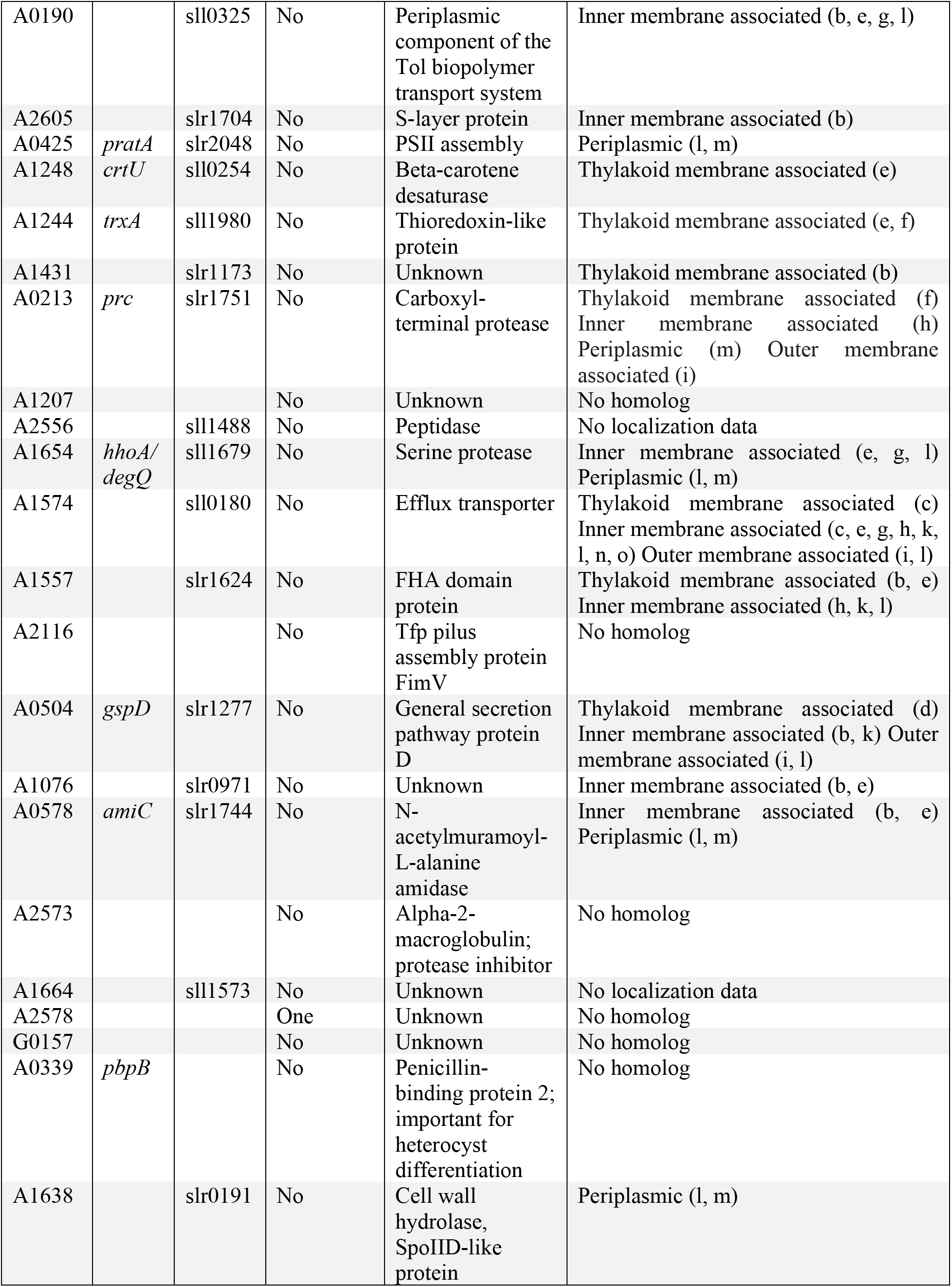
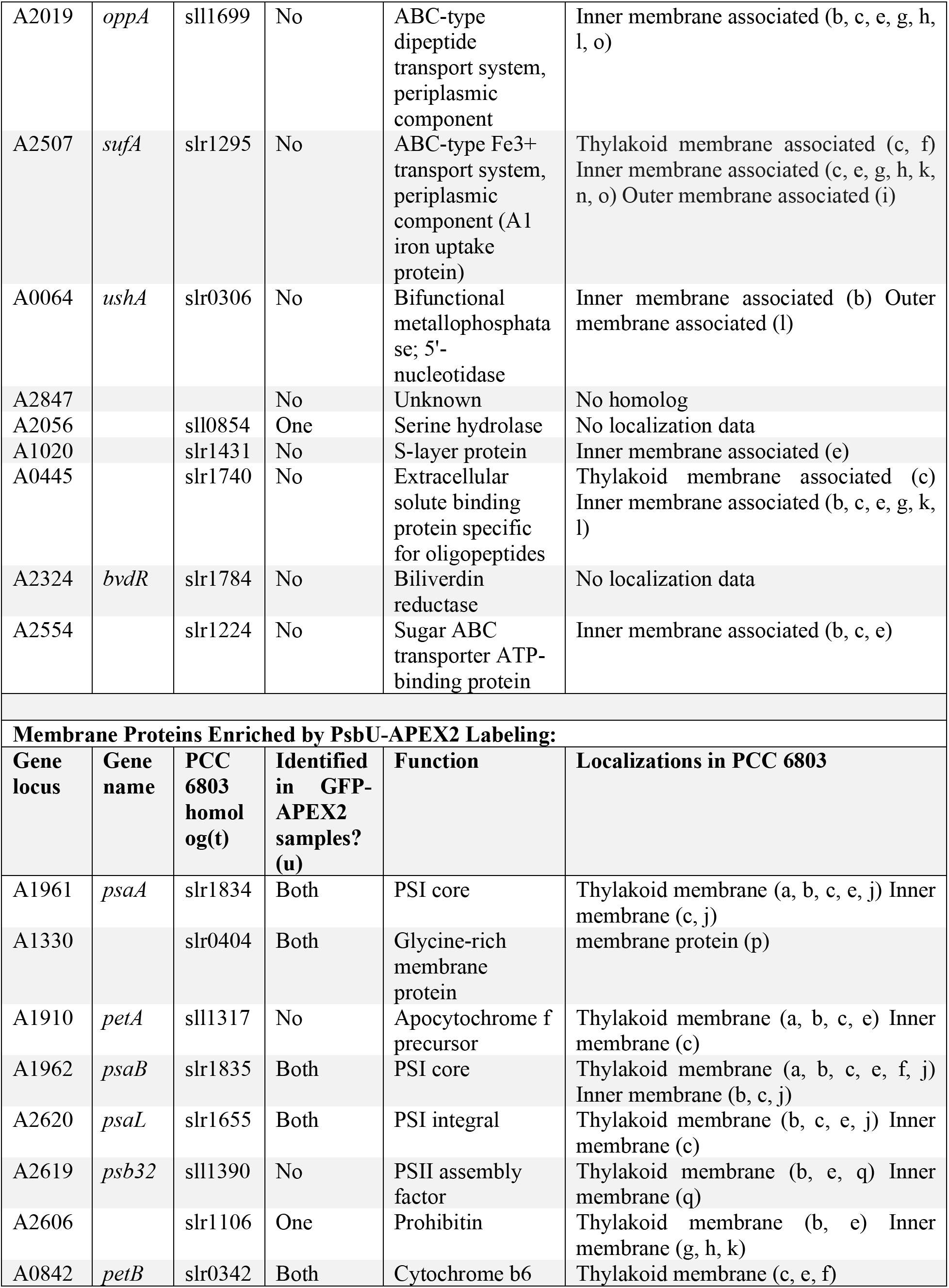
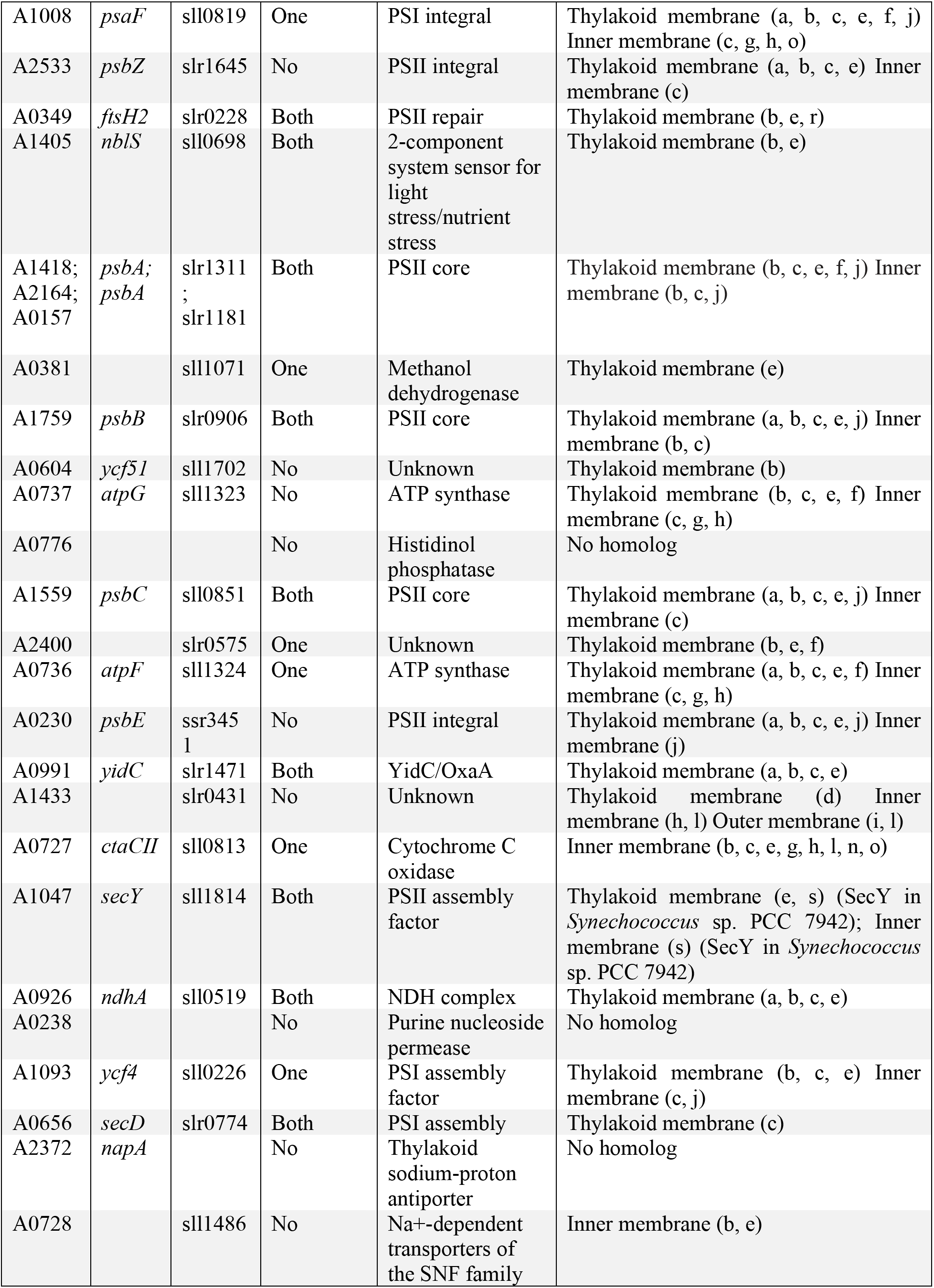
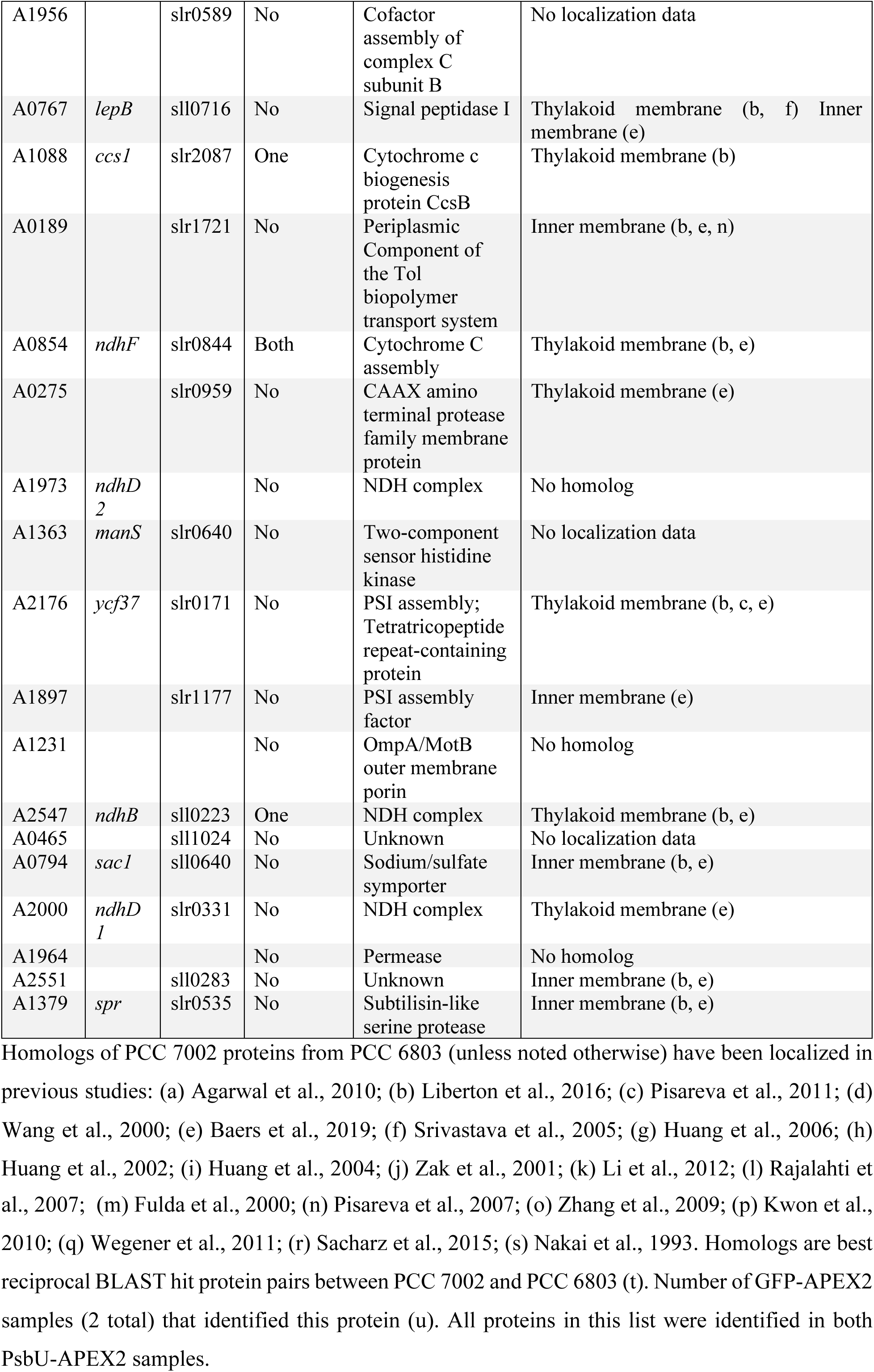
Proteins Enriched by PsbU-APEX2

The list of PsbU-APEX2 enriched proteins includes many proteins expected to be present within the thylakoid lumen. For example, the extrinsic subunits of the PSII water oxidizing complex (PsbU, PsbQ, PsbO, and PsbV) and Cyt *C*_6_ (PetJ1) were enriched by PsbU-APEX2. Many integral membrane proteins from PSII, photosystem I (PSI), and cytochrome b_6_f were also enriched. Furthermore, several factors involved in PSII and PSI assembly and PSII repair were also enriched (PsbP, Ycf48, CtpA, PratA, Psb32, FtsH2, YidC, SecY, Ycf4, SecD, and A1897). Interestingly, PSI extrinsic subunits (PsaC and PsaD) on the cytoplasmic side of PSI were also enriched, although these proteins were also identified in the cytoplasmic GFP-APEX2 samples. Several ATP synthase subunits were also enriched in the PsbU-APEX2 samples. In addition to proteins involved in photosynthetic electron transport, many proteins involved in cellular respiration, metabolite transport, redox regulation, and protein trafficking, processing, and assembly were enriched in PsbU-APEX2 samples.

Many proteins enriched in PsbU-APEX2 samples have previously been found in the periplasm or inner membrane (Table 1). It is unclear what biological relevance this may have. It is possibly an artifact of overexpression of PsbU-APEX2. However, the cyanobacterium *Gloeobacter violaceus* does not contain a thylakoid membrane (Mareš et al., 2013) and instead performs oxygenic photosynthesis in the inner membrane. If the thylakoid membrane and lumen originated from the plasma membrane and periplasmic space, respectively, perhaps it is not surprising that some proteins are found in both cellular fractions. Furthermore, ultrastructural studies of *Synechocystis* sp. PCC 6803 using cryo-electron tomography identified sites of contact between the thylakoid and plasma membrane (Rast et al., 2019). Additional possibilities include dual localization of proteins, low fidelity of the sorting mechanism of translocated proteins into the lumen and the periplasm, and post-translocation sorting of proteins into their final localization. Further experiments are needed to determine the biological relevance of the periplasmic and inner membrane proteins observed. There were several proteins enriched by PsbU-APEX2 that have not been previously localized or have unknown functions. These proteins could be the subject of future research.

The experiments performed here demonstrate the potential of APEX2 to interrogate the proteome of regions of cyanobacteria that have not been previously biochemically purified, like the thylakoid lumen. In the future, this technique can be used to monitor the proteomes of other regions of the cell under different environmental conditions. Additionally, APEX2 can be used to determine the topology of membrane proteins and identify candidates for protein-protein interactions (Lee et al., 2016; Lobingier et al., 2017; Mavylutov et al., 2018; Paek et al., 2017). Proximity-based proteomics using APEX2 has the potential to be a powerful tool in the pursuit of understanding the physiology of photosynthetic organisms.

## Methods

### Creation of PCC 7002 Strains

The *psbU* gene (SynPCC7002_A0322) was amplified from PCC 7002 while APEX2 was amplified from a plasmid gifted to us by Alice Ting (Addgene plasmid # 72558; http://n2t.net/addgene:72558; RRID:Addgene_72558). Plasmids were assembled using Gibson Assembly (Gibson et al., 2009) with NS1 as the homology arms, *pccmK2* as the promoter (Cameron et al., 2013), and kanamycin resistance for selection. The Gibson reactions were transformed into DH5α *E. coli,* and minipreps of liquid cultures started from single colonies were performed to collect plasmid. Plasmid was transformed into PCC 7002 (Stevens and Porter, 1980) and colonies containing the desired insert were serially passaged in the presence of antibiotic until segregated.

### Biotinylation of Proteins by APEX2 in PCC 7002

Biotinylation of proteins was performed using a modified protocol from Hung et al. and Hwang and Espenshade (2016; 2016). Briefly, 50 mL cultures of PCC 7002 strains were grown in A+ media (Stevens et al., 1973) in air at 37°C with a light intensity of 185 μmol photons • m^-2^ • s^-1^ for 2 days to an OD730 of about 0.5. Several μL of culture were saved to image on the microscope. The culture was pelleted at 4300 x *g* for 10 minutes at 4°C. The supernatant was poured off and cells were resuspended in 4 mL A+ with 2.5 mM BP and transferred to a six-well plate. Six-well plates were incubated shaking in air at 37°C with a light intensity of 185 μmol photons • m^-2^ • s^-1^ for 30 minutes. Samples were then pelleted in a 1.5 mL tube and resuspended in 1 mL phosphate buffered saline pH 7.8 (Bio-Rad) (PBS). 10 μL of 100 mM H_2_O_2_ was added and cells were inverted for 30 seconds before pelleting for 30 s. Supernatant was removed and cells were resuspended in quencher solution (PBS with 10 mM sodium ascorbate, 5 mM Trolox and 10 mM sodium azide) and pelleted. This step was repeated two additional times. The supernatant was removed and the cell pellets were frozen at −80°C for storage and to facilitate cell lysis.

### Cell Lysis

The cell pellet was resuspended in RIPA lysis buffer with quenchers (50 mM Tris pH 7.4, 150 mM NaCl, 0.1% (w/v) SDS, 0.5% (w/v) sodium deoxycholate, 1% (v/v) Triton X-100, 10 mM sodium ascorbate, 5 mM Trolox, 10 mM sodium azide, 1 mM PMSF). Cells were lysed using bead beating, with 30 cycles of 20 seconds on and 20 seconds off on ice. The lysate and beads were pelleted at 5000 x g in a microfuge and the supernatant was collected. The supernatant was clarified to remove debris and unbroken cells following centrifugation at 21,100 x g for 5 min at 4°C.

### Protein Concentration Measurement

The protein concentration of cell lysate was quantified using the Pierce 660 nm Protein Assay (Thermo Fisher).

### Purification of Biotinylated Proteins

Streptavidin magnetic beads (Pierce) were washed twice in RIPA lysis buffer (50 mM Tris pH 7.4, 150 mM NaCl, 0.1% (w/v) SDS, 0.5% (w/v) sodium deoxycholate, 1% (v/v) Triton X-100) and the supernatant was removed. 800 μL of RIPA lysis buffer with quenchers containing 50 μg of protein for every 50 μL of streptavidin magnetic beads was added. Beads were incubated with protein for 1 hour at room temperature on a rotator. The beads were then washed twice with RIPA lysis buffer, once with 1M KCl, once with 0.1 M Na2CO3, once with 8 M urea in 10 mM Tris pH 7.5, and once again with RIPA lysis buffer.

### Elution of Biotinylated Proteins for gels and blots

Beads were boiled in 30 μL of elution buffer (3X Laemmli buffer, 2 mM biotin, 20 mM DTT) to elute biotinylated proteins. The eluate was collected and diluted with 60 μL of water to run on gels.

### Preparation for Mass Spectrometry

Beads were washed an additional 5 times with 50 mM NH4HCO3 containing 0.2% (w/v) sodium deoxycholate. The supernatant was removed and beads were resuspended in 50 μL 10 mM TCEP and 40 mM chloroacetamide and incubated at 37°C for 30 minutes to reduce and alkylate the proteins. 150 μL water containing 0.225% (w/v) sodium deoxycholate and 0.2 μg Promega sequencing grade modified trypsin was added. An on-bead digestion was performed overnight on a rotator at 37°C. Beads were pelleted and the supernatant was collected. Formic acid was added to 2% (w/v) to stop digestion. Sodium deoxycholate was removed using 3 phase transfers with ethyl acetate. The samples were desalted using in-house STAGE tips with 3M Empore SDB-RPS membrane and dried using a vacuum centrifugation.

### LC-MS/MS

The tryptic peptides were resolved using an UltiMate 3000 UHPLC system (Thermo Fisher) in a direct injection mode. Peptides were reconstituted in Buffer A (0.1% formic acid in water), and peptide concentration was measured using Fluoraldehyde o-Phthaldialdehyde Reagent (Thermo Fisher). For each sample, 250 ng (5 μL) of the peptides were loaded onto a Waters BEH C18 column (130 Å, 1.7 μm × 75 μm × 250 mm) with 98.4% Buffer A and 1.6% Buffer B (0.1% formic acid in acetonitrile) at 0.4 μL/min for 16.67 min. Peptides were resolved and eluted using a gradient of 1.6 to 8% B (0-8 min), 8-20% B (8-140 min), and 20-32% B (140-160 min) at 0.3 μL/min. MS/MS was performed on a Q-Exactive HF-X mass spectrometer (Thermo Fisher), scanning precursor ions between 380-1580 m/z (60,000 resolution, 3 x 10^6^ ions AGC target, 45 msec maximum ion fill time), and selecting the 12 most intense ions for MS/MS (15,000 resolution, 1 x 10^5^ ions AGC target, 150 ms maximum ion fill time, 1.4 m/z isolation window, 27 NCE, 30 s dynamic exclusion). Ions with unassigned charge state, +1, and > +7 were excluded from the MS/MS.

### Silver Stain Protocol

Proteins were separated on a 10% SDS-PAGE gel and stained using the short silver nitrate staining protocol described in by Chevallet et al. (2006).

### Immunoblotting

Proteins were separated on a 10% SDS-PAGE gel and immunoblots were performed following the protocol from Green and Sambrook (2012). Protein was transferred to a nitrocellulose membrane, or a polyvinylidene fluoride (PVDF) membrane if fluorescent secondary antibodies were used. After blocking membranes overnight, membranes were incubated with GFP (Invitrogen, cat. no. A6455) or RbcL (Agrisera, cat. no. AS03037) antibodies, or streptavidin-HRP (Life Technologies, cat. no. R960-25). Membranes probed for GFP or RbcL were then incubated with a secondary antibody conjugated to HRP or AlexaFluor 488 (Thermo Fisher, cat. no. A-11008 or cat. no. 31460). Membranes were visualized using chemiluminescence after exposure to the Clarity Western ECL substrate (Bio-Rad) or fluorescence. If necessary, blots were stripped using ReBlot Plus Mild Solution (Millipore).

### Fluorescence Microscopy

Cells were spotted onto an agar pad (A+ with 1% agar) and placed onto a microscope slide. Cells were imaged on a customized Nikon TiE inverted wide-field microscope with a Near-IR-based Perfect Focus system. Images were acquired with an ORCA Flash4.0 V2+ Digital sCMOS camera (Hamamatsu) using a Nikon CF160 Plan Apochromat Lambda 100x oil immersion objective (1.45 N.A.). Chlorophyll fluorescence of thylakoid membranes was imaged using a 640 nm LED light source (SpectraX) for excitation and a standard Cy5 emission filter. GFP localization was imaged using a 470 nm LED light source (SpectraX) for excitation and a standard GFP emission filter.

### LC-MS/MS Data Analysis

Only proteins with at least two unique peptides and two spectral counts were considered identified. Proteins identified in both GFP-APEX2 replicates and/or both PsbU-APEX2 replicates were retained for further analysis. A presence/absence Venn diagram was constructed (Figure 3C). A protein must be identified in both replicates of a sample to appear in the Venn diagram. Proteins identified in both replicates of a sample and only one replicate of the other sample (176 proteins) were not added to the Venn diagram as their localization was unclear.

The log_2_ ratio of the MAXQUANT LFQ intensities and the log_2_ ratio of normalized spectral counts were used as metrics to determine enrichment in the PsbU-APEX2 A and PsbU-APEX2 B samples over the GFP-APEX2 B sample (log_2_(U/G)) (Old et al., 2005). If a protein was not identified in a sample, the LFQ intensity was set to zero. To determine the cutoff for proteins enriched in PsbU-APEX2 samples, identified proteins were cross-referenced with true positive (TP) or false positive (FP) lists. The TP lists were assembled using localization data from proteomic studies in PCC 6803 (Agarwal et al., 2010; Fulda et al., 2000; Huang et al., 2002, 2004, 2006; Li et al., 2012; Liberton et al., 2016; Pisareva et al., 2007, 2011; Rajalahti et al., 2007; Srivastava et al., 2005; Wang et al., 2000; Zak et al., 2001; Zhang et al., 2009). The FP list contained proteins annotated as DNA binding or involved in transcription or translation by McClure et al. (2016). TP proteins included all thylakoid lumen protein homologs from Rajalahti et al. (2007). PCC 7002 homologs of proteins found associated with the thylakoid membrane by Pisareva et al. (2011) and at least one other study were included in the TP list, otherwise TP protein homologs were found in at least three thylakoid membrane studies. TP proteins were required to possess a secretion signal or at least one transmembrane helix to be retained in the TP list and had to be found in equal to or more thylakoid proteomics studies than inner membrane/outer membrane/periplasm studies (Supplemental Table 3).

A total of four analyses were performed, one for each enrichment metric in each PsbU-APEX2 sample. For each protein in every analysis, the true positive rate (TPR) and the false positive rate (FPR) were calculated using only the proteins with enrichment equal to or greater than the protein of interest. The TPR was the number of TP proteins found at or above the cutoff divided by the total number of TP proteins found in the experiment. The FPR was the number of FP proteins found at or above the cutoff divided by the total number of FP proteins. The cutoff for each sample was the enrichment with the greatest TPR-FPR value. The proteins above the cut-off of the in all 4 analyses are reported in Table 1.

### Signal Sequence Prediction

To predict if a protein had a signal sequence and the cut site to the remove the signal sequence, all proteins in the UniProt reference proteome for PCC 7002 were analyzed with SignalP-5.0 using both the Gram-positive and Gram-negative bacterial options.

### Transmembrane Helices Prediction

To predict if a protein had transmembrane helices, all proteins in the UniProt reference proteome for PCC 7002 were analyzed using the TMHMM Server v. 2.0.

## Author Contributions

K.K.D., T.L., and J.C.C. designed experiments. K.K.D. and T.L. performed experiments. K.K.D. and C.G. analyzed data. K.K.D. and J.C.C. wrote the manuscript.

## Acknowledgements

We would like to thank Patrick Thomas for the original design of plasmid constructs and Kristin A. Moore for critical analysis of manuscript. We thank all members of the Cameron laboratory for helpful discussions when designing experiments and analyzing data. This material is based upon work supported in part by the U.S. Department of Energy, Office of Science, Office of Biological and Environmental Research under Award Number DE-SC0019306 to J.C.C. This work was supported in part by the Interdisciplinary Quantitative Biology (IQ Biology) program at the BioFrontiers Institute, University of Colorado, Boulder, NSF IGERT grant number 1144807. Financial support for this study was also provided by startup funds from the University of Colorado-Boulder to J.C.C. The authors declare no conflicts of interests.

## SUPPLEMENTAL INFORMATION

**This manuscript contains the following supplemental information:**

**Supplemental Data Analysis**

**Supplemental Figure 1.** Comparison of enrichment values between samples.

**Supplemental Figure 2.** Comparison of FP and TP enrichment values.

**Supplemental Figure 3.** ROC curves and enrichment cutoffs.

**Supplemental Table 1.** Proteomics data and analysis

**Supplemental Table 2.** Additional information about enriched proteins

**Supplemental Table 3. List of true and false positive proteins for analysis**

### Supplemental analysis of LC/MS-MS data

After calculating the enrichment metrics for each analyzed protein, these values were plotted in histograms to check for normal distributions (Supplemental Figure 1A-D). Furthermore, the enrichment ratios from each sample for the same sample were compared against each other to check for correlation between samples (Supplemental Figure 1E-F). Pearson’s correlation coefficient was calculated for the enrichment values of samples A and B using both LFQ intensities and spectral counts. For both the LFQ intensities and spectral counts, the enrichment values had a Pearson’s correlation coefficient of 0.96, indicating a strong positive correlation between samples.

Next, box plots were made comparing the enrichment distribution of all proteins, TP proteins, and FP proteins for each sample and each enrichment metric (Supplemental Figure 2A-D). In each comparison, the enrichment of TP proteins had a greater mean than the FP proteins. A student t-test found significant (p<0.5) differences between the enrichment values of TP and FP proteins in both enrichment metrics in each sample, supporting the fact that the experiment did enrich for proteins biotinylated by PsbU-APEX2 localized in the thylakoid lumen.

After confirming that thylakoid lumen and thylakoid membrane proteins were enriched in the experiments, Receiver Operating Curve (ROC) were plotted for each sample. For this analysis, proteins were organized in descending enrichment value for both enrichment metrics in each sample (Supplemental Figure 3A-D). The True Positive Rate (TPR), was calculated for each protein as the % of all identified TP proteins with an enrichment greater than or equal to itself. Conversely, the False Positive Rate (FPR) was calculated for each protein as the % of all identified FP proteins with an enrichment greater than or equal to itself. In ROCs, the TPR is plotted against FPR for each individual protein and the graph is examined to determine if the line arcs over a TPR = FPR line, which demonstrates that TPs are enriched in the experiment. For each enrichment metric in each sample, the ROC curve arced over the TPR = FPR line, indicating that TP proteins were enriched in the experiment. Furthermore, the cutoff for enriched proteins was placed at the enrichment value were the TPR-FPR was at a maximum. To visualize this, the TPR - FPR was plotted against enrichment in each sample (Supplemental Figure 3E-H).

**Supplemental Figure 1.**
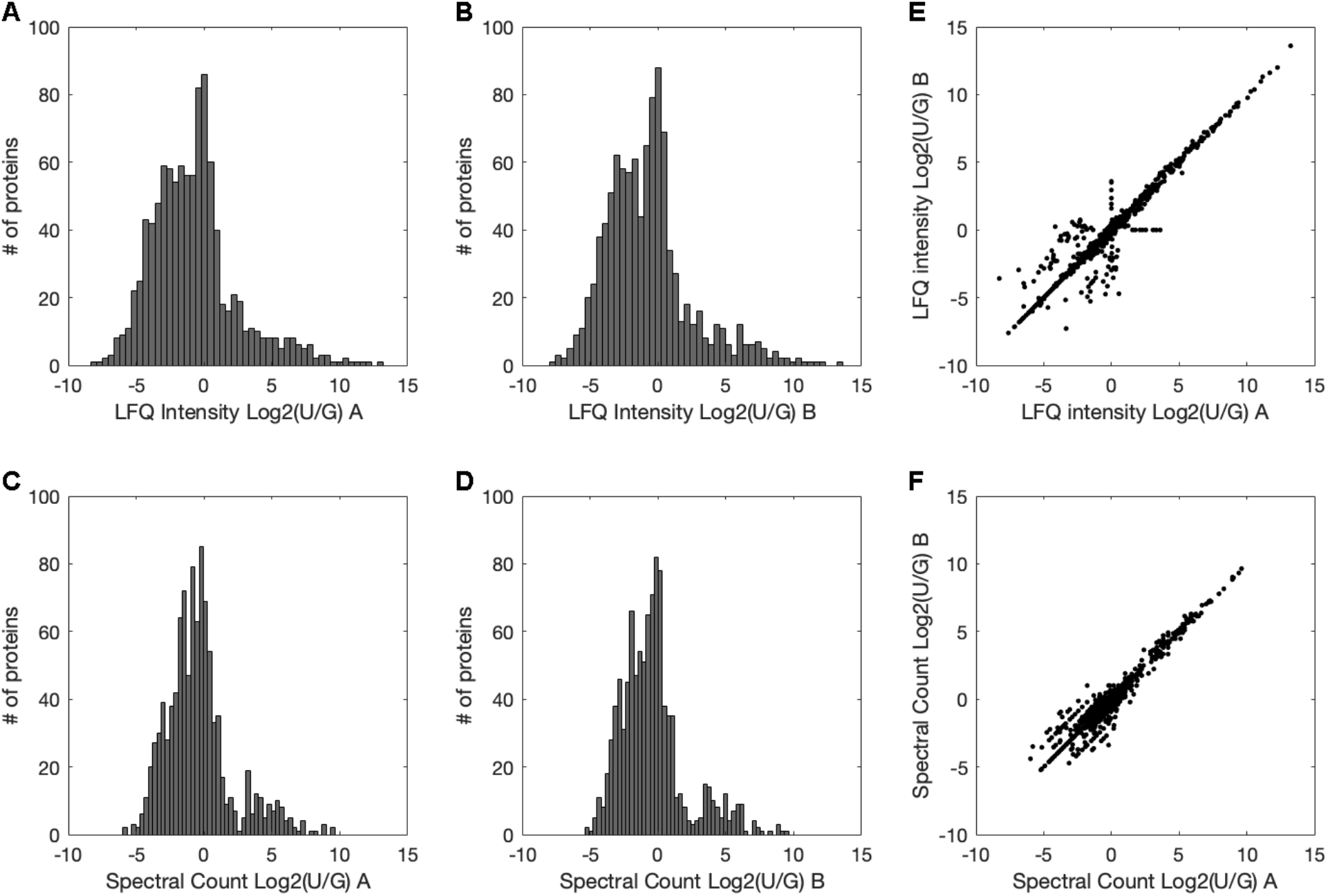
Comparison of enrichment values between samples. **(A)** Histogram of enrichment using LFQ intensities from PsbU-APEX2 sample A. **(B)** Histogram of enrichment using LFQ intensities from PsbU-APEX2 sample B. **(C)** Histogram of enrichment using spectral counts from PsbU-APEX2 sample A. **(D)** Histogram of enrichment using spectral counts from PsbU-APEX2 sample B. **(E)** Comparison of LFQ intensity enrichment values for each analyzed protein between PsbU-APEX2 samples A and B. PCC is 0.96. **(F)** Comparison of spectral count enrichment values for each analyzed protein between PsbU-APEX2 samples A and B. PCC is 0.96.

**Supplemental Figure 2.**
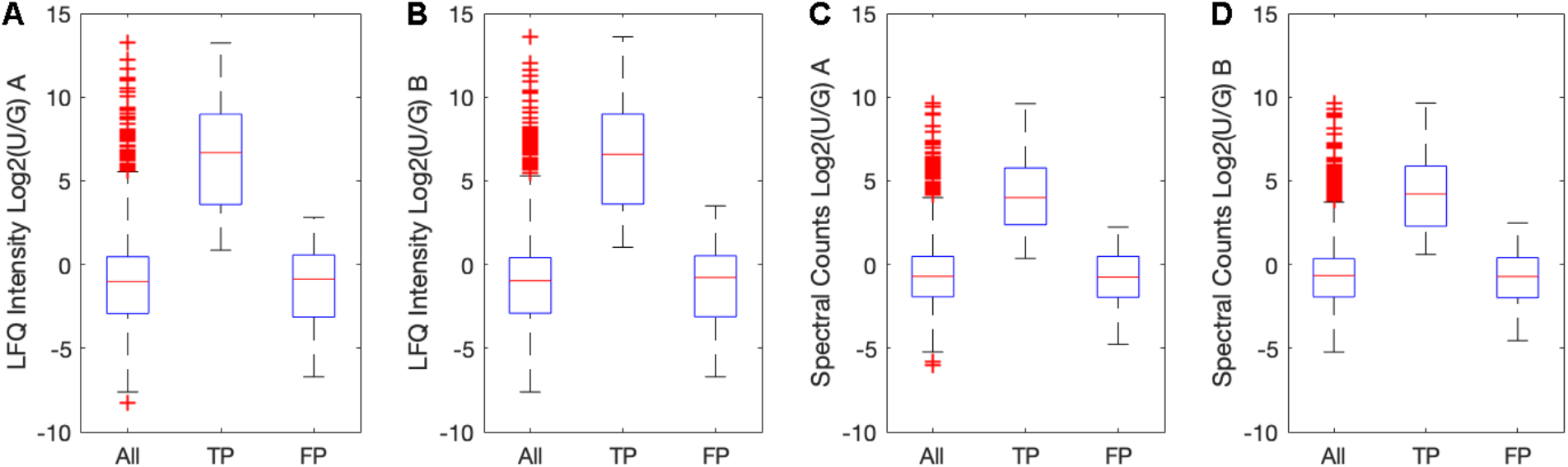
Comparison of FP and TP enrichment values. (A) Distributions of enrichment values calculated using LFQ intensity in sample A. (B) Distributions of enrichment values calculated using LFQ intensity in sample B. (C) Distributions of enrichment values calculated using spectral counts in sample A. (D) Distributions of enrichment values calculated using spectral counts in sample B. All panels show the distribution enrichment values for all proteins, True Positive (TP) proteins, and False Positive (FP) proteins in a sample in box and whisker plots. In each sample the TP enrichment values are shifted up compared to the FP enrichment values. A Student’s t-test found significant differences (p < 0.05) between the TP and FP proteins in each sample (A, p = 4 x 10^-15^; B, p = 7 x 10^-15^; C, p = 6 x 10^-14^; D, p = 5 x 10^-14^). Red crosses signify outliers, which are greater than 75^th^ percentile + 2.97 • standard deviation • (75^th^ percentile-25^th^ percentile) or less than 25^th^ percentile −2.97 • standard deviation • (75^th^ percentile-25^th^ percentile).

**Supplemental Figure 3.**
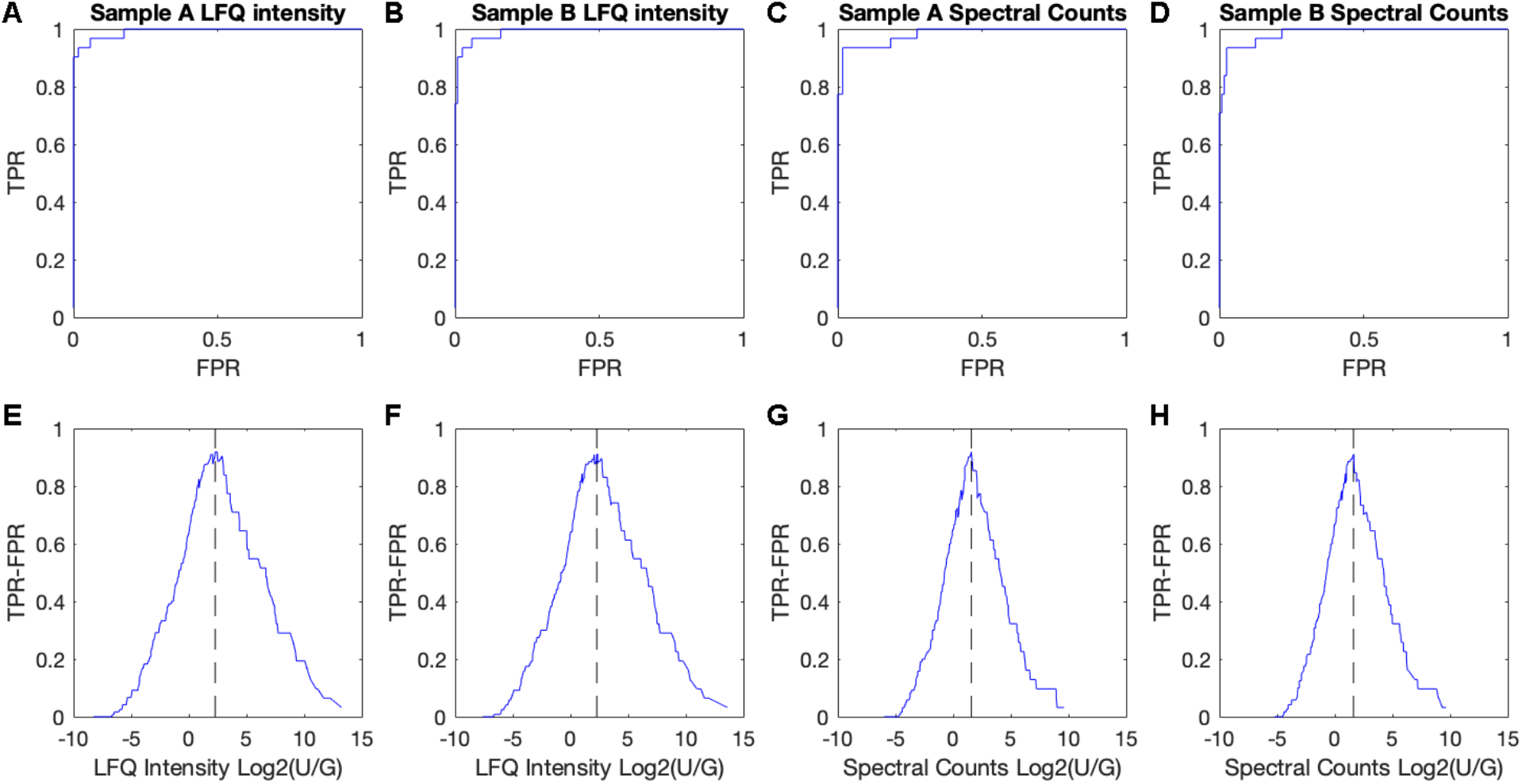
ROC curves and enrichment cutoffs. The top row of samples shows ROC curves **(A-D)**, which plot TPR vs FPR to determine if the curve arcs over the TPR=FPR line. In all cases the ROC arcs over the TPR=FPR line, indicating that TP proteins are enriched in the experiment. The bottom row of samples shows TPR-FPR vs enrichment metrics **(E-H)**. The cutoff was drawn at the enrichment with the greatest TPR-FPR and is indicated by the dashed line. **(A)** ROC curve made using the enrichment values calculated using LFQ intensity in sample A. **(B)** ROC curve made using the enrichment values calculated using LFQ intensity in sample B. **(C)** ROC curve made using the enrichment values calculated using spectral counts in sample A. **(D)** ROC curve made using the enrichment values calculated using spectral counts in sample B. **(E)** TPR-FPR vs enrichment in sample A calculated using LFQ intensity. Cutoff is at 2.28. **(F)** TPR-FPR vs enrichment in sample B calculated using LFQ intensity. Cutoff is at 2.28. **(G)** TPR-FPR vs enrichment in sample A calculated using spectral counts. Cutoff is at 1.58. **(H)** TPR-FPR vs enrichment in sample B calculated using spectral counts. Cutoff is at 1.62.

